# Binding to m^6^A RNA promotes YTHDF2-mediated phase separation

**DOI:** 10.1101/757658

**Authors:** Jiahua Wang, Liyong Wang, Jianbo Diao, Yujiang Geno Shi, Yang Shi, Honghui Ma, Hongjie Shen

**Affiliations:** Center for Medical Research and Innovation, Shanghai Pudong Hospital, Fudan University Pudong Medical Center, and Institutes of Biomedical Sciences, Fudan University, 2800 Gongwei Road, Pudong, Shanghai 201399, China; Division of Newborn Medicine and Epigenetics Program, Department of Medicine, Boston Children’s Hospital, Boston, MA 02115, USA; Cell Biology Department, Harvard Medical School, Boston, MA 02115, USA; Key Laboratory of Arrhythmias of the Ministry of Education of China, East Hospital, Tongji University School of Medicine, Shanghai 200120, China; Division of Endocrinology, Diabetes and Hypertension, Department of Medicine, Brigham and Women’s Hospital, Harvard Medical School, Boston, MA, USA

## Abstract

As the most abundant modification on mRNA in mammal, N6-Methyladenosine (m^6^A) has been demonstrated to play important roles in various biological processes including mRNA splicing, translation and degradation. m6A reader proteins have been shown to play central roles in these processes. One of the m6A readers, YTHDF2 is localized to the P granules, which are liquid-like droplets where RNA degradation occurs. How YTHDF2 is localized to P granules is unknown. Here we provide evidence that YTHDF2 forms liquid droplets and phase separate, mediated by its low complexity (LC) domains. Interestingly, the ability to phase separate is robustly stimulated by m^6^A RNAs in vitro. In vivo, YTHDF2 phase separation may in fact be dependent on m^6^A RNA and YTHDF2 binding to m^6^A RNA, since a YTHDF2 m^6^A-binding defective mutant or a wildtype YTHDF2 assayed in cells lacking m^6^A RNAs, both fail to phase separate. The ability of phase separate is not limited to YTHDF2; we find other members of the YTH-domain m^6^A readers can also undergo phase separation. Our findings suggest that m^6^A RNA induced phase separation of m^6^A readers may play an important role in their distributions to different phase-separated compartments in cells.

## Main

As one of the most abundant modifications on mRNA in mammal, N^6^-methyladenosine (m^6^A) has been demonstrated to play important roles in various biological processes including nuclear RNA export, mRNA splicing, miRNA processing, mRNA degradation and translation(Shi et al.,2019). Importantly, different m^6^A reader proteins have been shown to play central roles in these processes.

YTH (YT521-B homology)-domain containing proteins are members of the conserved m^6^A reader family, which recognize m^6^A via the YTH domain(Hazra et al.,2019). Members of this family have been shown to play a role in mRNA translation and mRNA decay(Wang et al.,2014; Wang et al.,2015; Li et al.,2017; Shi et al.,2017). However, mechanisms by which these readers impact various biological processes are still elusive. For instance, YTHDF2 has been shown to regulate mRNA decay and is localized to the membrane-less cytoplasmic P granules(Wang et al.,2014), where mRNA decay occurs. But how YTHDF2 and its associated m^6^A RNAs are localized to this phase-separated granule(Standart and Weil,2018) remains largely unknown.

As P bodies are liquid-like droplets in cytoplasm (Standart and Weil,2018), we speculated that YTHDF2 may share liquid-like phase separation (LLPS) features. As with many LLPS proteins, YTHDF2 contains low a complexity (LC) domain (aa 230-383), which includes a glutamine (Q) rich domain (aa 288-383) (Figure 1A). Interestingly, recombinant YTHDF2 protein containing the LC domain (YTHDF2^aa 230-383^) forms liquid droplets (23 μM protein in 37 mM NaCl, 10% PEG8000), which are sensitive to 1,6-hexanediol (previously shown to specifically disrupt liquid-like assemblies)(Kroschwald et al.,2017) (Figure 1B), suggesting phase separation. This phase separation ability is dependent on NaCl concentration (Supplementary Figure 1A), which is consistent with another report(Ries et al.,2019). Moreover, we found these droplets fuse with each other to form bigger droplets, which further suggests a liquid like phase separation feature of YTHDF2^aa 230-383^ (Supplementary Figure 1B, and supplementary Video 1). Other members of this family, YTHDF1 and YTHDF3, also contain glutamine (Q) rich domain (Supplementary 1C), and also have phase separation ability (Supplementary 1D). As YTHDF2 glutamine compositional bias is conserved among vertebrates, to determine whether glutamine richness within the LC domain is important for LLPS of YTHDF2, we changed all glutamine to alanine in YTHDF2^aa 230-383^ (Supplementary Figure 1E), and the mutated protein essentially failed to form as many and large LLPS under the same assay condition (Supplementary Figure 1F, 1G). Similarly, another phase separation protein MED1 contains conserved serine (S) rich region, and this serine bias is necessary for MED1 phase separation(Sabari et al.,2018)(Sabari et al.,2018). To test whether YTHDF2 forms LLPS in cell, we overexpressed EGFP-YTHDF2 in mouse embryonic stem (ES) cells and human U2OS cells. FRAP assays showed YTHDF2 forms LLPS in both cell lines (Figure 1C, 1D).

**Figure 1.**
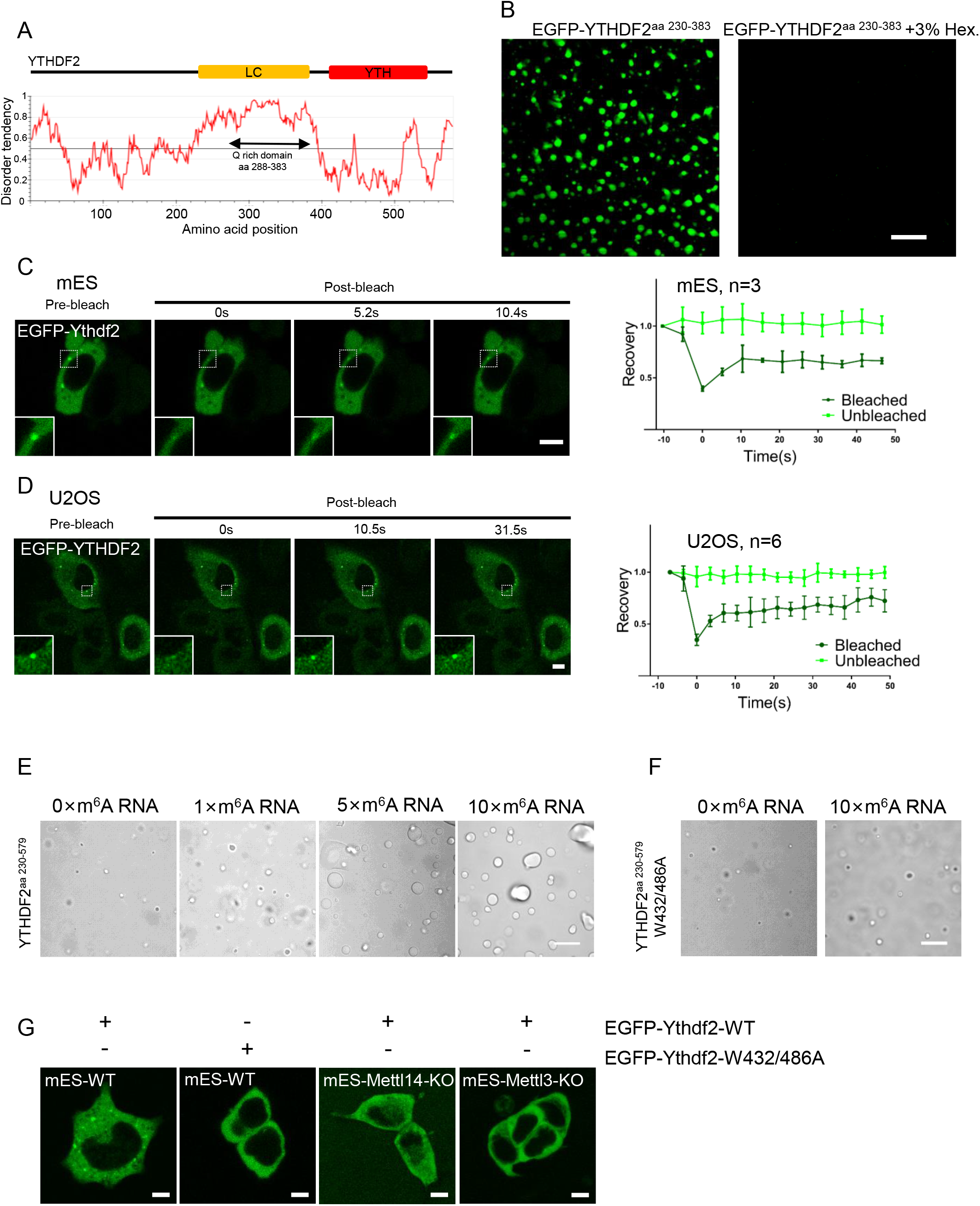
Phase separation of YTHDF2 in vivo/vitro. (A) Top: Diagram of protein domains of YTHDF2. Bottom: predictions of Intrinsic disorder tendency of YTHDF2 by IUPred2A(https://iupred2a.elte.hu/). Scores above 0.5 indicate disorder. (B) Liquid phase separation of YTHDF2^aa 230-383^ (23 μM YTHDF2, 37 mM NaCl, 10% PEG8000) was sensitive to 1,6-hexanediol (1,6-hex; 3%). Scale bar, 10 μm. (C, D) EGFP-YTHDF2 was exogenously expressed in mES cells (C) or U2OS cells (D). FRAP assays showed YTHDF2 forms LLPS in both cell lines (Left). The line traces represent mean fractional fluorescence (Right). Scale bar, 5 μ (C), 10 μm (D). (E) m^6^A oligos induced liquid like droplet formation of YTHDF2^aa 230-579^ (58 μM YTHDF2, 33 mM NaCl, 17 μM RNA oligos). Scale bar, 10 μm. (F) m^6^A oligos could not induce liquid like droplet formation of W432A/W486A mutated YTHDF2^aa 230-579^ (58 μM YTHDF2, 33 mM NaCl, 17 μM RNA oligos). Scale bar, 10 μm. (G) Wildtype, but not W432A/W486A mutated EGFP-YTHDF2 formed droplets in mESCs. Wildtype EGFP-YTHDF2 failed to form droplets in Mettl14 or Mettl3 KO mESCs. Scale bar, 5 μm.

Previous studies suggested that RNA plays a crucial role in protein phase separation. For instance, RNA has been shown to facilitate HP1 alpha phase separation on heterochromatin(Strom et al.,2017). Given that YTHDF2 preferentially binds m^6^A RNA(Wang et al.,2014), we hypothesized that m^6^A RNA might promote YTHDF2-mediated phase separation. To test this idea, we constructed a portion of the YTHDF2 protein containing aa 230 to its C terminus (aa 579), which includes both the LC and the YTH domains (Figure 1A). We identified a condition (13 μM YTHDF2, 37 mM NaCl, 0.74 μM RNA, 10% PEG8000) under which the recombinant YTHDF2^aa 230-579^ protein barely formed LLPS with RNA containing RNA oligos containing no m^6^A in vitro (Supplementary Figure 1H, right and 1I). Importantly, however, under the same assay condition, RNA oligos containing one m^6^A modification induced liquid like droplet formation (Supplementary Figure 1H, left and 1I). In addition, we found this phase separation enhancement appears to be dependent on the number of m^6^A in the RNA. As shown in Figure 1E, phase separation mediated by YTHDF2 increases in a manner that is dependent on the number of m6A sites in the RNA oligos (50bp RNAs containing either 0, 1, 5, or 10 m^6^A). These results are consistent with the recent reports that RNA oligos containing multiple m^6^A methylation sites robustly induce droplet formation by YTHDF2(Gao et al.,2019; Ries et al.,2019). Consistently, induction of phase separation of YTHDF2 by m6A RNAs seems to be dependent on the YTH domain, as even the RNA containing 10 m^6^A failed to enhance phase separation in vitro when the m6A-binding capability of the YTH domain is compromised (YTHDF2^aa 230-579^ carrying W432A, W486A)(Li et al.,2014) (Figure 1F). To corroborate this finding, we investigated whether binding of YTHDF2 to m^6^A is necessary for phase separation in vivo by expressing wildtype and the m^6^A-binding defective YTHDF2 (YTH mutated (W432A, W486A)) in mouse ES cells. While wildtype YTHDF2 formed droplets in cell, YTH mutated YTHDF2 did not (Figure 1G), indicating that the ability of YTHDF2 to bind m^6^A RNA is necessary for YTHDF2 LLPS in vivo. To further confirm this finding, we asked whether YTHDF2 forms LLPS in cells lacking the mRNA m^6^A enzymatic complex, METTL3 or METTL14 (Supplementary Figure 1J). We detected no YTHDF2 droplets in these cells (Figure 1G), further supporting the notion that m^6^A promotes YTHDF2 LLPS in cells.

In summary, we provide both in vitro and in vivo data demonstrating that m^6^A enhances phase separation of YTHDF2. Although recombinant YTHDF2 itself can phase separate in vitro, m^6^A modification significantly enhances this ability, and in vivo, YTHDF2 LLPS may in fact be dependent on binding m^6^A mRNAs.

While this manuscript was in preparation, several groups reported that YTHDF family proteins display phase separation potential(Fu and Zhuang,2019; Gao et al.,2019; Ries et al.,2019) stimulated by m^6^A RNA(Gao et al.,2019; Ries et al.,2019), consistent with our finding that m^6^A promotes the phase separation potential of YTHDF2.

---

**Supplementary Figure 1.**
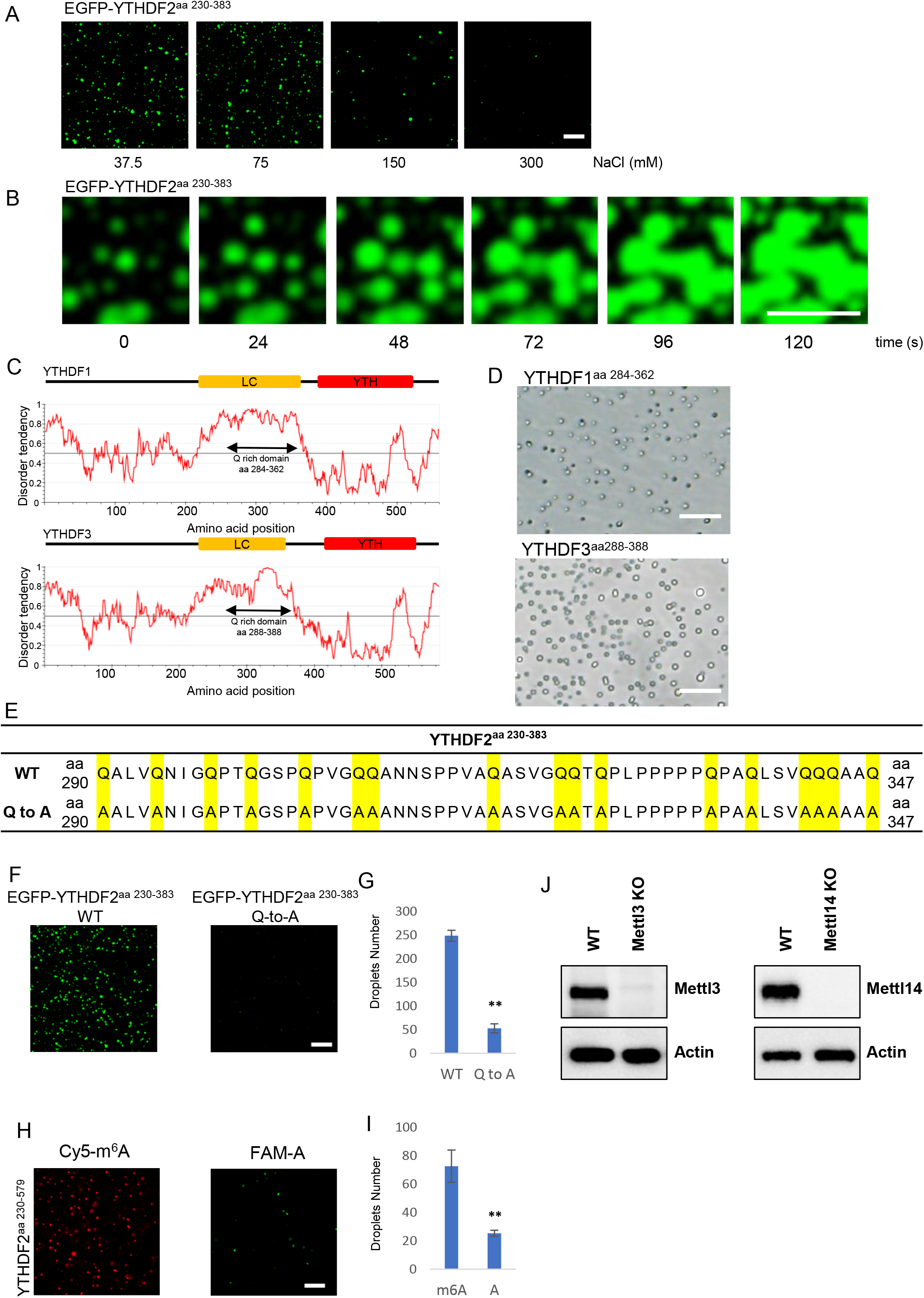
(A) Microscopy images of EGFP-YTHDF2^aa 230-383^ droplets (23 μM YTHDF2, 10% PEG8000) subjected to a NaCl gradient, showing NaCl concentration affects the formation of protein droplets. Scale bar, 10 μm. (B) The fusion of YTHDF2^aa 230-383^ droplets (23 μM YTHDF2, 37 mM NaCl, 10% PEG8000) were imaged by microscopy over 120 seconds. Scale bar, 10 μm. (C) Predictions of Intrinsic disorder tendency of YTHDF1 and YTHDF3 by IUPred2A(https://iupred2a.elte.hu/). Scores above 0.5 indicate disorder. (D) Liquid phase separation of YTHDF1^aa 284-362^ (Glutamine rich domain) and YTHDF3^aa 288-388^ (Glutamine rich domain). Scale bar, 10 μm. (E) Sequence alignment of the wildtype and Q-to-A mutant of the Glutamine (Q) rich domain in YTHDF2^aa 230-383^. (F) Q-to-A mutation compromised liquid phase separation of YTHDF2^aa 230-383^ (11 μM YTHDF2, 37 mM NaCl, 10% PEG8000). Scale bar, 10 μm. (G) Droplet numbers of wildtype and Q-to-A mutated YTHDF2^aa 230-383^ proteins. Data are represented as mean from three replicates. Error bars represent SEM of the number of droplets from three replicates. **P < 0.01, t test. (H) Cy5-m^6^A, but not FAM-A RNA oligos induced YTHDF2^aa 230-579^ liquid like droplet formation (13 μM YTHDF2, 37 mM NaCl, 0.74μM RNA, 10% PEG8000). (I) Droplet numbers of Cy5-m^6^A and FAM-A RNA oligos induced YTHDF2^aa 230-579^ proteins. Data are represented as mean from three replicates. Error bars represent SEM of the number of droplets from three replicates. **P < 0.01, t test. (J) Western blot showing Mettl3 protein levels in WT, Mettl3 KO and Mettl14 protein levels in WT, Mettl14 KO cell lines.

## AUTHOR CONTRIBUTIONS

J.W. and L.W. carried out all the experiments described in this manuscript. H.M. and Y.S. directed all the experiments with input from J.D. H.S. and Y.S. conceived the project and co-wrote the manuscript.

## ACKNOWLEDGEMENTS

H.S. was sponsored by Shanghai Rising-Star Program (19QA1401300) and National Science Foundation of China (81874157, 31601060). Research was also supported by funds from Fudan University. YS is an American Cancer Society Research Professor.

## MATERIALS AND METHODS

### Prediction of Intrinsically Unstructured Proteins

Predictions of intrinsic disorder tendency of the proteins were accomplished by specifying the accession numbers of the proteins in the “Enter SWISS-PROT/TrEMBL identifier or accession number” field of IUPred2A(Meszaros et al., 2018) (https://iupred2a.elte.hu/). The red line in the result page represents the intrinsic disorder tendency of each protein.

### Cell cultures

U2OS cells were cultured in Dulbecco’s Modified Eagle’s Medium (DMEM) supplemented with 10% fetal bovine serum (FBS) and 100 U/mL Penicillin/Streptomycin at 37 °C with 5% CO2. E14Tg2a murine embryonic stem cells were cultured in DMEM supplemented with 10% FBS, 1% MEM nonessential amino acid, 55 mM β-Mercaptoethanol, 1000 U/mL LIF (Millipore) and 100 U/mL Penicillin/Streptomycin at 37 °C with 5% CO2.

### Stable cell lines construction

EGFP-tagged murine Ythdf2 was cloned into the pPB-CAG-IRES-Pac plasmid with N-terminal Flag and HA tags. This plasmid was individually co-transfected with pCMV-PBase plasmid into mESCs using Lipofectamine 2000 (Invitrogen) according to the manufacturer’s instruction. Medium was replaced by fresh media with 2 μg/mL Puromycin after 48 hours. After continuous Puromycin selection for 3 days, the survived mESCs were pooled as stable overexpression cell lines.

EGFP-tagged human YTHDF2 was cloned into the pLenti6.2-V5 plasmid. Lentivirus was made by co-transfection of this vector with VSV-G and psPAX2 in a 3:1:1 ratio into 293T cells. Supernatant at 48 hours post-transfection was collected and concentrated by PEG8000. U2OS cells were seeded in a 6-well plate and infected with lentivirus supernatant in the presence of 5 μg/mL polybrene (Sigma). Medium was replaced by fresh media with 10 μg/mL Blasticidin S at 24 hours post-infection. After continuous Blasticidin S selection for 5 days, survived U2OS cells were pooled as stable infected cell lines.

CRISPR-Cas9 gene targeting was carried out as previously described(Maeder et al., 2013) and the single knockout clones were isolated and then confirmed by Western blot showing undetectable Mettl3 and Mettl14 protein. Guiding RNA sequences used are: 1) Mettl3 KO: 5’-GCTTAGGGCCGCTAGAGGTA-3’. 2) Mettl14 KO: 5’-GTAGCTCAGCAGGTGTGCGG-3’.

### Protein Expression and Purification

The different truncated fragments of human YTHDF2 were subcloned into the pMCSG7 vector with the sequence EGFP-GSGS (linker) or not. The YTHDF2-LC Q-to-A mutant DNA was synthesized and inserted into the same vector. All proteins were expressed in Rosetta (DE3) cells. Cells in 500 ml LB media were induced overnight at 16 °C with 0.1mM Isopropyl β-D-Thiogalactoside(IPTG) and collected. The pellets were resuspended in 30 ml lysis buffer (50 mM Tris pH7.5, 500 mM NaCl, 10 mM imidazole, 0.01% NP40, 1mM β-Mercaptoethanol) and squeezed three times at 4 °C. After centrifugation at 13000 rpm, 4 °C for 20 minutes, the supernatants were purified with Ni-NTA beads (Smart-Lifesciences) and eluted by elution buffer (50 mM Tris pH7.5, 500 mM NaCl, 200 mM imidazole, 1mM β-Mercaptoethanol). Elutions containing proteins were then analyzed by Coomassie staining and dialyzed against storage buffer (50 mM Tris pH7.5, 100 mM NaCl, 1 mM β-Mercaptoethanol).

### In vitro droplet assay

Purified proteins were concentrated to the indicated protein concentration described in the text using Amicon Ultra centrifugal filters (10K MWCO, Millipore) and added to solutions at varying concentrations with the indic ated final salt and molecular crowder concentrations. The droplet assay was performed in the following buffer: 50 mM Tris (PH7.5), 1mM β-Merc aptoethanol, and indicated NaCl. The droplet assay was generated by co mbining YTHDF2 protein or YTHDF2/RNA mixture with 40% PEG8000 in a ratio of 3:1, if PEG8000 was used. The protein solutions were loaded onto a confocal dish and imaged with 10%PEG8000 (Figure 1B, S1A, S1B, S1F, S1H, S1J) or not (Figure 1E, 1F). For detail, in m^6^A-RNA induc ed droplet assays, probe and protein were firstly mixed in a microtube a nd imaged on a confocal dish. The sequences of the different m^6^A-RNA probes are 5’-Cy5-CGUGG m^6^ACUGGCU-3’ or 5’-FAM-CGUGGACUGGC U-3’. The sequences of the different sites m^6^A-RNA probes are: 1) 0x m ^6^A-RNA: GGACUGGACUGGACUGGACUGGACUGGACUGGACUGGACUGGACUGGACU. 2) 1x m^6^A-RNA: GGACUGGACUGGACUGGACUGGm^6^AC UGGACUGGACUGGACUGGACUGGACU. 3) 5x m^6^A-RNA: GGm^6^ACUGG ACUGGm^6^ACUGGACUGGm^6^ACUGGACUGGm^6^ACUGGACUGGm^6^ACUGG ACU. 4) 10x m^6^A-RNA: GGm^6^ACUGGm^6^ACUGGm^6^ACUGGm^6^ACUGGm^6^A CUGGm^6^ACUGGm^6^ACUGGm^6^ACUGGm^6^ACUGGm^6^ACU.

### Super-resolution microscopy

Stable cell lines were seeded in 35mm confocal dishes and imaged with a STED Nanoscopes leica TCS SP8 STED. The images were post-processed by the Leica LAS AF Lite software.

### Fluorescence recovery after photobleaching (FRAP)

The FRAP was performed on the STED Nanoscopes leica TCS SP8 STED with 488 nm laser. Bleaching was performed over an approximately 1 μm^2^ region using 100% laser power and images were collected every 3.5 seconds (U2OS) or 5.2 seconds (mESC). Fluorescence intensity was measured using the LAS X software. Background intensity was subtracted and values were reported relative to pre-bleaching time points.

